# Sendai virus persistence questions the transient naive reprogramming method for iPSC generation

**DOI:** 10.1101/2024.03.07.583804

**Authors:** Alejandro De Los Angeles, Clemens B. Hug, Vadim N. Gladyshev, George M. Church, Sergiy Velychko

## Abstract

Since the revolutionary discovery of induced pluripotent stem cells (iPSCs) by Shinya Yamanaka, the comparison between iPSCs and embryonic stem cells (ESCs) has revealed significant differences in their epigenetic states and developmental potential. A recent compelling study published in *Nature* by Buckberry et al.^1^ demonstrated that a transient-naive-treatment (TNT) could facilitate epigenetic reprogramming and improve the developmental potential of human iPSCs (hiPSCs). However, the study characterized bulk hiPSCs instead of isolating clonal lines and overlooked the persistent expression of Sendai virus carrying exogenous Yamanaka factors. Our analyses revealed that Sendai genes were expressed in most control PSC samples, including hESCs, which were not intentionally infected. The highest levels of Sendai expression were detected in samples continuously treated with naive media, where it led to overexpression of exogenous MYC, SOX2, and KLF4, altering both the expression levels and ratios of reprogramming factors. Our findings call for further research to verify the effectiveness of the TNT method in the context of delivery methods that ensure prompt elimination of exogenous factors, leading to the generation of bona fide transgene-independent iPSCs.

## Detection of Sendai virus sequences in established pluripotent stem cell lines

Our analysis of publicly available RNA-seq data provided by Buckberry et al. revealed significant expression of Sendai virus genes in nearly all PSC samples (**Fig. 1a**). The highest levels of Sendai genes were observed in naive hiPSCs, suggesting a possible selection of cells with persistent viral expression^2,3^. Similar findings were published by Yamanaka and colleagues reporting the persistence of Cytotune 2.0 Sendai viruses in human naive PSCs cultured in t2iLGo conditions^4^. One of the two primed hiPSC samples showed expression of Sendai virus genes as late as passage 17. Traces of Sendai virus were also found in naive-to-primed (NTP) hiPSCs, which were established and passaged in naive media before being transferred into primed PSC conditions, whereas TNT hiPSCs were Sendai-free. Surprisingly, control MEL1 hESC samples from Buckberry et al. also had low but detectable levels of Sendai expression. In contrast, hESC samples from another group^5^ did not express any Sendai genes, as expected. hESCs were not deliberately infected with Sendai viruses, suggesting a contamination or possibly a mix-up of hESC and TNT hiPSC samples.

**Fig. 1:**
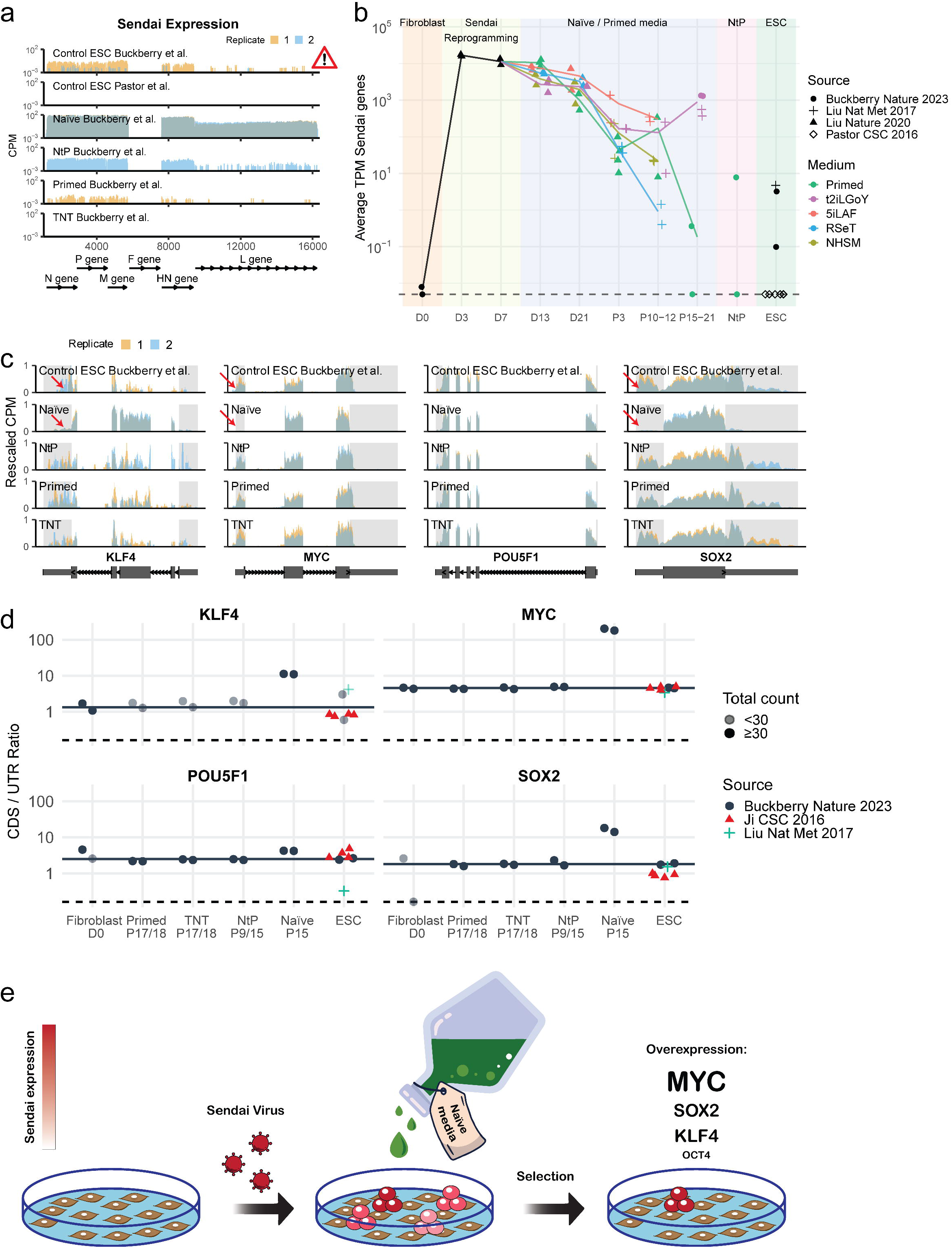
Persistence of Sendai virus and selection for exogenous factor expression by naive media. **a**, Read coverage of the Sendai virus genome. Two replicates per condition are shown (in red and blue). Coverage was binned at 100 bp resolution and is shown as counts per million reads (CPM). The F gene was deleted in the commercial kit used for reprogramming in these studies. **b**, Time-course of Sendai expression across reprogramming and hPSC samples. The y-axis shows the mean expression as transcripts per million (TPM) of all Sendai genes – except for F. Symbols indicate the study from which samples were sourced and colors indicate the cell culture media that the samples were grown in. The dashed line at the bottom corresponds to the location of samples with zero TPM. **c**, Read coverage of Yamanaka factors. The CPM values are rescaled such that each sample has a range from exactly zero to one. Large boxes in the gene models indicate exons, small boxes correspond to 5’ and 3’ untranslated regions (UTRs), and lines with arrows indicate introns. UTRs are highlighted using gray shading. **d**, Quantification of the ratio between coding sequence and untranslated region expression (CDS/UTR ratio). For each Yamanaka factor we quantified the number of reads aligning to CDS and UTR regions. The ratio of normalized counts CDS/UTR serves as a proxy for the proportion of transcripts originating from the viral transgene vs the endogenous locus. The horizontal black line is drawn at the mean ratio of the hESC control samples from Buckberry et al., which serve as baseline for the expected ratio in the absence of a transgene. The horizontal dashed line is drawn at the location of samples with a ratio of zero. Samples with low expression below <30 raw counts are drawn in a lighter shade. **e**, Graphical overview and mechanism of TNT method: the study utilizes t2iLGoY naive media, which preferentially selects for cells expressing high levels of exogenous reprogramming factors from Sendai virus RNA (illustrated in dark red). This selective expression of exogenous factors may significantly contribute to the outcomes observed in the TNT reprogramming.

## Naive media selects for high levels of exogenous Yamanaka factors

We assessed the levels of Sendai virus sequences in RNA-seq data across human reprogramming intermediates and established naive and primed hiPSC bulk lines, encompassing Buckberry et al. and two additional studies by the same group (Liu et al., Nature Methods 2017; and Liu et al., Nature 2020)^1,6,7^ (**Fig. 1b**). The analyses confirmed the absence of Sendai virus in control fibroblast samples as well as expected pronounced expression levels in Day 3 and Day 7 reprogramming samples. For each of the culture conditions tested (t2iLGoY, 5iLAF, NHSM, RSeT—naive medias, and E8—primed media), the levels of Sendai viruses decreased with passaging, yet remained detectable even at late passage numbers. The presence of Sendai virus in control hESCs from Liu et al, 2017 was also noted. Finally, a nearly 1,000-fold higher level of Sendai expression was observed in t2iLGoY-cultivated hiPSCs relative to primed hiPSCs at similar high passage numbers, which amounts to approximately two-fold enrichment for Sendai-expressing cells during each passage in naive media (every 3 days).

A significant presence of Sendai virus sequences in control naive hiPSCs enabled us to evaluate the expression level of each Yamanaka factor. Endogenous transcripts include both coding sequences (CDS) and untranslated regions (UTRs), while the Sendai RNA contains only the CDS sequences allowing discrimination of the endogenous and exogenous RNAs. Examining the RNA-seq read coverage for the Yamanaka factors revealed a striking difference in expression patterns between naive cells and other samples: naive samples showed significantly lower UTR expression while maintaining high CDS coverage, suggesting that most transcripts are derived from loci lacking UTRs (**Fig. 1c**). To quantify the extent of exogenous expression, we evaluated the ratio of reads mapping to CDSs versus UTRs. Our analysis found higher CDS/UTR ratios for MYC, SOX2, and KLF4 (SKM) in naive hiPSCs compared to the expected CDS/UTR ratio found in samples with low or absent transgene expression, such as fibroblasts and hESCs, indicating a selection for exogenous SKM expression in the naive hiPSCs (**Fig. 1d-e**). Exogenous MYC was the most enriched, with a CDS/UTR ratio greater than 100, followed by SOX2 and KLF4 with ratios exceeding 10. Exogenous OCT4, however, was not noticeably enriched, which likely reflects relatively high levels of endogenous OCT4 expression in primed PSCs.

The resulting overexpression of SKM in reprogramming samples treated with naive media echoes our studies showing that omitting OCT4 from the Yamanaka cocktail could improve the quality of iPSCs^8^ and reset PSCs across species^9^, hinting at the possible mechanism underlying TNT reprogramming.

## Implications and recommendations for future studies

Prior research emphasized the need for transgene clearance to ensure reactivation of the endogenous pluripotency network and to avoid issues like abnormal differentiation. Even minimal leakage of exogenous OSKM from a tet-inducible promoter compromises the developmental potential of iPSCs, rendering tet-inducible OSKM iPSCs incapable of producing healthy animals^8-9^. Our results suggest that the t2iLGoY naive medium, used in the TNT reprogramming protocol, promotes the retention of exogenous reprogramming factors and changes their ratios (**Fig. 1e**). This raises concerns about applications of such media in reprogramming protocols. Yamanaka factors can induce a transient naive-like state even in primed media^10^. Identification of the optimal reprogramming factor ratio could eliminate the need for the naive medium treatment with its associated risks of genetic and epigenetic instability^11^. Refinements of the Cytotune 2.0 Sendai kit, such as an all-in-one design or the addition of microRNA-binding sites, might promote efficient virus elimination. Meticulous assessment for exogenous factor elimination is crucial. Furthermore, we recommend isolating and characterizing clonal iPSC lines rather than bulk passaging of reprogramming cultures, to reduce heterogeneity and the potential for selection.

## Conclusion

The study by Buckberry et al. suggests that a specific naive media regimen can boost the hiPSC technology. Our finding of Sendai virus sequences in control hiPSC and hESC lines, coupled with naive media favoring high Sendai expression suggests that further work needs to be done to support the effectiveness of the TNT reprogramming method, underscoring the necessity for a refined delivery system.

## Methods

### Processing of RNA-seq data

Raw sequencing reads were downloaded from SRA accession numbers SRP286549 (Buckberry et al.^1^), SRP259918 (Liu et al. 2020^7^), SRP115256 (Liu et al. 2017^6^), SRP059279 (Ji et al. 2016^12^), and SRP068579 (Pastor et al. 2016^5^).

### RNA-seq alignment and quantification

Human transcripts were quantified using human genome version GRCh38 and version 110 of the Ensembl gene annotations. For Sendai virus transcripts we used sequences and gene annotations from NCBI reference sequence NC_075392.1 (*Respirovirus muris*). We quantified transcript expression using Salmon^13^ 1.10.2 using the options `-l A --seqBias --gcBias --posBias --softclip`. The Salmon index was created by concatenating the CDS fasta file from Ensembl with the Sendai transcripts obtained from NC_075392.1. The raw transcript counts from Salmon were imported into R 4.3.1 using the tximport package^14^ and aggregated to gene-level counts. For each RNA-seq sample, a library specific correction factor accounting for differences in sequencing depth was calculated using `estimateSizeFactorsForMatrix()` from DESeq2^15^.

Additionally, all reads were aligned to the human genome using STAR^16^ 2.7.9a with default settings. The genome index was prepared by concatenating the unmasked primary DNA assembly from Ensembl with the whole Sendai genome.

### Read coverage plots

Read coverage plots for the Sendai genome and Yamanaka factors were generated based on our STAR alignments using the ggcoverage R package. Reads were counted in evenly spaced 100 bp bins, normalized using the previously calculated size factors, divided by the mean number of reads per sample, and then multiplied by a million to get Counts Per Million (CPM). For comparing coverage of coding sequences (CDS) to 5’ and 3’ untranslated regions (UTR) we rescaled raw CPM values for each sample so that their minimum and maximum are zero and one, respectively.

### Calculation of CDS to UTR ratios

The ratio between expression of CDS to UTRs in naturally occurring mature mRNAs is expected to be close to one, given how they are usually transcribed and spliced as a single unit. mRNA transcribed from exogenous viral sequences do not contain UTRs, therefore shifting the CDS/UTR ratio up the more they are expressed. Read counts for the coding sequences (CDS), as well as 5’ and 3’ untranslated regions (UTR) for each Yamanaka factor were quantified based on our STAR alignments using the Rsubread^17^ R package. For each of the four factors we picked a representative transcript, annotated as “MANE select” in Ensembl. These were ENST00000325404, ENST00000259915, ENST00000621592, and ENST00000374672 for SOX2, POU5F1, MYC, and KLF4, respectively. Raw counts for each feature type and transcript were added up, normalized using DESeq2 size factors, and divided by the total feature length. These normalized counts for CDS and UTR for each transcript were then divided by each other to yield CDS/UTR ratios.

## Acknowledgements

We thank Caitlin MacCarthy for editing the manuscript. We acknowledge support from the NIA grant R01 AG058063 (C.H.) and NCI U54-CA225088 (C.H.).

## Author contributions

S.V. and A.A. conceived the study. A.A. performed initial analysis. C.H. performed an independent full analysis and generated the figures. A.A., S.V. and C.H. interpreted the results and wrote the manuscript.

V.N.G. and G.M.C. advised on the study.

## Competing interests

S.V. is listed as an inventor of a submitted patent on SK/SKM naive reset. All other authors declare no competing interests.

## References

1. Buckberry, S. et al. Transient naive reprogramming corrects hiPS cells functionally and epigenetically. Nature 620, 863–872 (2023).

2. Takashima, Y. et al. Resetting Transcription Factor Control Circuitry toward Ground-State Pluripotency in Human. Cell 158, 1254–1269 (2014).

3. Guo, G. et al. Naive Pluripotent Stem Cells Derived Directly from Isolated Cells of the Human Inner Cell Mass. Stem Cell Rep. 6, 437–446 (2016).

4. Kunitomi, A. et al. Improved Sendai viral system for reprogramming to naive pluripotency. Cell Rep. Methods 2, 100317 (2022).

5. Pastor, W. A. et al. Naive Human Pluripotent Cells Feature a Methylation Landscape Devoid of Blastocyst or Germline Memory. Cell Stem Cell 18, 323–329 (2016).

6. Liu, X. et al. Comprehensive characterization of distinct states of human naive pluripotency generated by reprogramming. Nat. Methods 14, 1055–1062 (2017).

7. Liu, X. et al. Reprogramming roadmap reveals route to human induced trophoblast stem cells. Nature 586, 101–107 (2020).

8. Velychko, S. et al. Excluding Oct4 from Yamanaka Cocktail Unleashes the Developmental Potential of iPSCs. Cell Stem Cell 25, 737-753.e4 (2019).

9. MacCarthy, C. M. et al. Highly cooperative chimeric super-SOX induces naive pluripotency across species. Cell Stem Cell 31, 127-147.e9 (2024).

10. Cacchiarelli, D. et al. Integrative Analyses of Human Reprogramming Reveal Dynamic Nature of Induced Pluripotency. Cell 162, 412–424 (2015).

11. Bar, S., Schachter, M., Eldar-Geva, T. & Benvenisty, N. Large-Scale Analysis of Loss of Imprinting in Human Pluripotent Stem Cells. Cell Rep. 19, 957–968 (2017).

12. Ji, X. et al. 3D Chromosome Regulatory Landscape of Human Pluripotent Cells. Cell Stem Cell 18, 262–275 (2016).

13. Patro, R., Duggal, G., Love, M. I., Irizarry, R. A. & Kingsford, C. Salmon provides fast and bias-aware quantification of transcript expression. Nat. Methods 14, 417–419 (2017).

14. Soneson, C., Love, M. & Robinson, M. Differential analyses for RNA-seq: transcript-level estimates improve gene-level inferences [version 2; peer review: 2 approved]. F1000Research 4, (2016).

15. Love, M. I., Huber, W. & Anders, S. Moderated estimation of fold change and dispersion for RNA-seq data with DESeq2. Genome Biol. 15, 1–21 (2014).

16. Dobin, A. et al. STAR: ultrafast universal RNA-seq aligner. Bioinformatics 29, 15–21 (2013).

17. Liao, Y., Smyth, G. K. & Shi, W. The R package Rsubread is easier, faster, cheaper and better for alignment and quantification of RNA sequencing reads. Nucleic Acids Res. 47, e47 (2019).

